# Integrated phenomenology and brain connectivity demonstrate changes in nonlinear processing in jhana advanced meditation

**DOI:** 10.1101/2024.11.29.626048

**Authors:** Ruby M. Potash, Sean D. van Mil, Mar Estarellas, Andres Canales-Johnson, Matthew D. Sacchet

**Affiliations:** Meditation Research Program, Department of Psychiatry, Massachusetts General Hospital, Harvard Medical School, Boston, MA, USA; Conscious Brain Lab, Department of Psychology, University of Amsterdam, Nieuwe Achtergracht 129-B, 1018 WT, Amsterdam, The Netherlands; Department of Biological and Experimental Psychology, Queen Mary University of London, United Kingdom; Department of Psychology, University of Cambridge, CB2 3EB Cambridge, United Kingdom; Neuroscience Center, Helsinki Institute of Life Science, University of Helsinki, P.O. Box 3, Fabianinkatu 33, FI-00014 Helsinki, Finland; Neuropsychology and Cognitive Neurosciences Research Center, Faculty of Health Sciences, Universidad Católica del Maule, 3460000 Talca, Chile

## Abstract

We present a neurophenomenological case study investigating distinct neural connectivity regimes during an advanced concentrative absorption meditation called jhana (ACAM-J),characterized by highly-stable attention and mental absorption. Using EEG recordings and phenomenological ratings (29 sessions) from a meditator with +20,000 hours of practice, we evaluated connectivity metrics tracking distinct large-scale neural interactions: nonlinear (WSMI and Directed Information), capturing non-oscillatory dynamics; and linear (WPLI) connectivity metrics, capturing oscillatory synchrony. Results demonstrate ACAM-J are better distinguished by non-oscillatory compared to oscillatory dynamics across multiple frequency ranges. Furthermore, combining attention-related phenomenological ratings with WSMI improves Bayesian decoding of ACAM-J compared to neural metrics alone. Crucially, deeper ACAM-J indicate an equalization of feedback and feedforward processes, suggesting a balance of internally- and externally-driven information processing. The results from this intensively sampled case study are a promising initial step in revealing the distinct neural dynamics during ACAM-J, offering insights into refined conscious states and highlighting the value of nonlinear neurophenomenological approaches to studying attentional states.

## INTRODUCTION

Meditation research is now expanding beyond mindfulness meditation into a new wave of study focused on advanced meditation (Sacchet et al., 2024). Advanced meditation includes states and stages of meditation that unfold with time and mastery. Advanced meditation includes transformative trajectories, that is, meditative development, which may culminate in outcomes known as meditative endpoints (Galante et al., 2023; Sparby and Sacchet, 2022; Wright et al., 2023). Meditative endpoints include recently neuroscientifically studied cessations of consciousness that lead to profound psychological insight and clarity (van Lutterveld et al., 2024a; Chowdhury et al., 2023; Laukkonen et al., 2023). Advanced concentrative absorption meditation (ACAM) is a class of advanced meditation that is characterized by self-induced states of highly stable attention and mental absorption. Jhana meditation (ACAM-J; jhāna in Pali, the liturgical language of Theravada Buddhism) is a type of ACAM that is characterized by a series of eight sequential states that are characterized by bliss. stability of mind, and expansiveness, and wherein spontaneous mental content is increasingly absent (Sparby and Sacchet, 2024; Yang et al., 2024b; Gunaratana, 1988; Sayadaw, 2008).

ACAM-J are associated with profound experiences of clarity, ego-dissolution, equanimity, and open-consciousness (Yang et al., 2024b; Sparby, 2019). During ACAM-J, practitioners systematically relinquish the dominant qualities of each of the states, allowing them to access increasingly deeper levels of ACAM-J. The meditator enters ACAM-J through an initial access concentration (AC) state, in which they focus their attention on a meditation object (e.g., breath, bodily feelings, visual imagery). The first four ACAM-J (ACAM-J1 to ACAM-J4) are classified as the “form” ACAM-J, while the later four (ACAM-J5 to ACAM-J8) are considered the “formless” ACAM-J. As meditators advance through the formless ACAM-J, consciousness, and perception become increasingly refined (Catherine, 2008; Rasmussen and Snyder, 2009; Sayadaw, 2008). For more details on the phenomenology and dominant qualities of the eight ACAM-J, refer to Sparby and Sacchet 2024; Yang et al. 2024b.

With (1) detailed historical documentation, (2) a systematic approach to practice, and (3) consistent phenomenology, ACAM-J is an ideal model for studying deep aspects of human consciousness; attention, narrative, and reward processing; and sensory attenuation (Yang et al., 2024b). Nonetheless, due to the rarity of ACAM-J practitioners and the necessary rigor of a neuroscientific methodology to meaningfully capture these nuanced meditative states, there have been limited studies of ACAM-J (Wright et al., 2023). Initial neuroimaging studies of ACAM-J point to a possible deconstruction of consciousness (Yang et al., 2024a; Treves et al., in press) with good within-subject reliability of functional MRI (fMRI) brain responses (Ganesan et al., 2024).

A distinctive phenomenological feature of ACAM-J is their marked change in the nature of attention. During ACAM-J, the meditator is absorbed with highly refined attention, often with no influence from external stimuli (Yang et al., 2024b). These features make ACAM-J a suitable model for investigating the neural dynamics of attention and perception. Insights from network neuroscience have demonstrated how meditation broadly influences brain networks, including shifts in connectivity patterns within key hubs (Prakash et al., in press). However, the mechanisms of large-scale communication of internally-driven perception are still debated. For example, it is debated whether attention relates to changes in linear connectivity manifested as oscillatory synchronization, or nonlinear connectivity manifested as non-oscillatory fluctuations in electrophysiological signals such as EEG (electroencephalography) and ECoG (electrocorticography) (Vinck et al., 2023; Spyropoulos et al., 2024). While the mainstream view is that attention might modulate functional connectivity between oscillations at the same frequency (Fries, 2015) (Figure 1A,B), other perspectives emphasize the nonlinear nature of neural communication which may require non-oscillatory rather than oscillatory dynamics (Vinck et al., 2023; Singer, 2021). Attention, in this alternative view, modulates nonlinear, non-oscillatory connectivity as a consequence of interactions across frequency bands rather than interactions within the same frequency band, as is the case of linear connectivity (Figure 1A,C). This idea is based on evidence demonstrating that nonlinear interactions are functionally relevant for complex computations such as pattern extraction in neural networks, leading to the integration of executive functions (Vinck et al., 2023; Singer, 2021), and for the long-range communication of perceptual and predictive information (Vinck et al., 2023; Gelens et al., 2024; Canales-Johnson et al., 2020a,b; Roberts et al., 2025). Thus, here we test the hypothesis that nonlinear connectivity metrics should outperform linear connectivity ones in distinguishing self-reported attentional states during ACAM-J.

**Figure 1:**
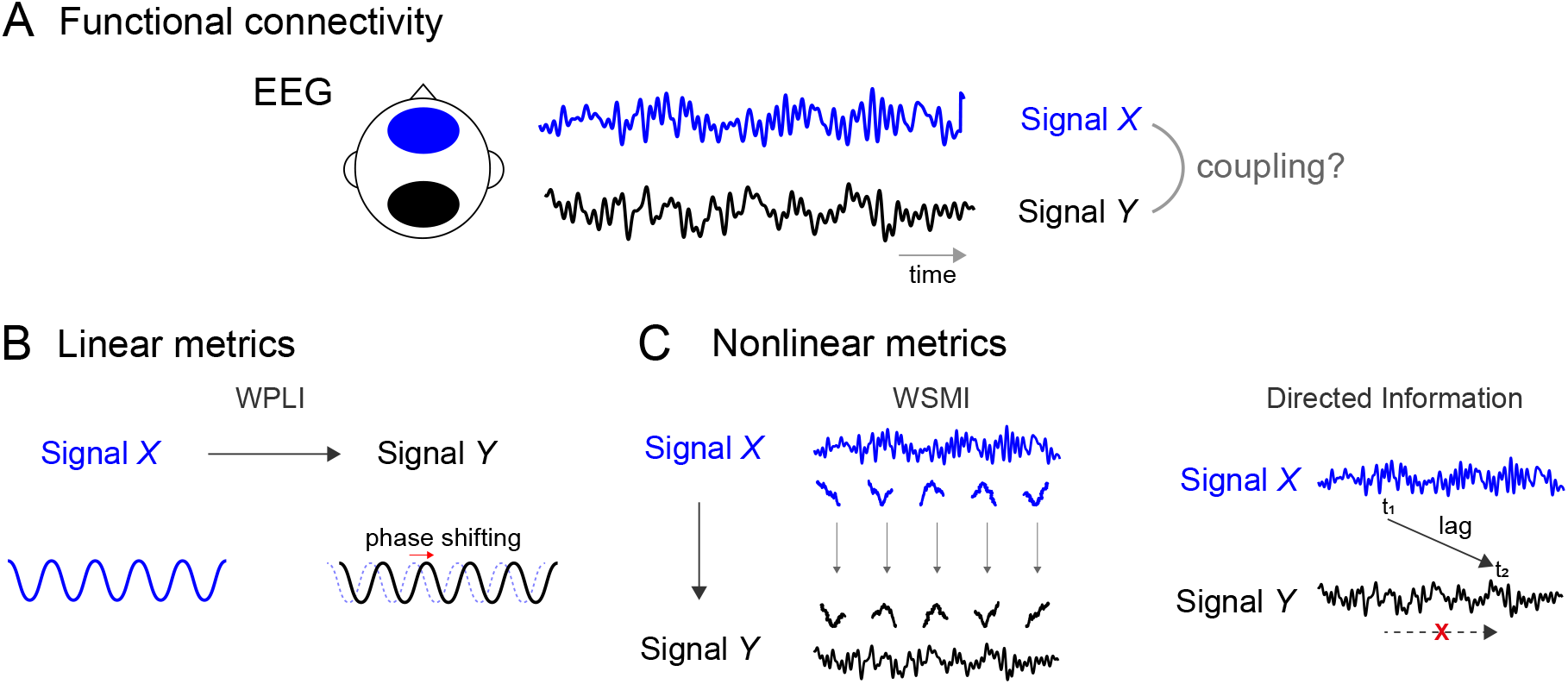
Experimental design, oscillatory and non-oscillatory connectivity. **(A)** Mapping functional connectivity between two EEG signals. In this diagram, connectivity is depicted between signal X (in blue) and signal Y (in black) corresponding to two arbitrary electrodes of interest. **(B)** Linear transformations between two correlated signals at the same frequency. The transformed signal (signal Y) is the result of a phase-shifted version of the original signal (signal X) at the same frequency (e.g., a 10 Hz signal X can result in a phase-shifted signal Y at the same 10 Hz). The WPLI metric (weighted phase lagged index) captures this type of linear, oscillatory connectivity. **(C)** Left: Nonlinear transformations cause systematic relationships across different frequencies between signal X and Y, potentially in the absence of linear transformations. Thus, non-linear connectivity measures quantify arbitrary mappings between temporal patterns in one signal (signal X) and another signal (signal Y). This type of non-oscillatory mapping is captured by the WSMI metric (weighted symbolic mutual information). Right: Directed Information (dir-INFO) (also referred to as Transfer Entropy) measures the coupling between the values of a signal Y, and the values of a signal X earlier in time, conditioning on the earlier values of Y itself. dir-INFO measures the time-lagged dependence between two signals, over and above the dependence with the past of the signal itself (its self-predictability).

In practice, nonlinear connectivity is captured from human electrophysiology using signal processing metrics derived from information theory, such as weighted symbolic mutual information (WSMI) (King et al., 2013; Imperatori et al., 2019) and Directed Information (dir-INFO) (Canales-Johnson et al., 2023; Ince et al., 2017) (Figure 1C), which show increased performance at distinguishing states and contents of consciousness than linear connectivity metrics. For example, WSMI outperforms linear connectivity metrics such as weighted phase lag index (WPLI) in distinguishing between sleep stages, healthy and pathological states of consciousness (King et al., 2013; Sitt et al., 2014; Imperatori et al., 2019; Canales-Johnson et al., 2020a), and changes in the contents of perception (Canales-Johnson et al., 2020b,a; Olivares et al., 2022). Additionally, dir-INFO, a nonlinear metric of directional functional connectivity, can distinguish between feedforward and feedback information flow in the human brain during perceptual tasks, providing insight into how sensory-generated information interacts with internally-generated neural dynamics during perception and cognition (Canales-Johnson et al., 2023). For example, directed information between frontal and posterior signals may be a mechanism for perceiving different contents during visual bistability (Canales-Johnson et al., 2023).

Here, we used EEG and phenomenological ratings collected during 29 sessions from a single subject with over 20,000 hours of meditation practice. We investigated changes in ACAM-J using a unique, intensive neurophenomenological case study approach that combines nonlinear neural dynamics, phenomenological ratings of attention, and classification techniques. Based on this intensive case study data, we showed that WSMI outperforms WPLI in distinguishing ACAM-J. Crucially, using a Naive Bayesian classifier for decoding, we found that integrating self-reported ratings of stability of attention improves the classification of ACAM-J when combined with nonlinear connectivity metrics (WSMI) compared to when we used this neural metric alone. Finally, we showed that deeper ACAM-J indicate a decrease in externally-driven information processing, quantified as a reduction in the brain’s feedforward compared to feedback information flow (dir-INFO). These findings lay the groundwork for future replication and validation in larger cohorts, further corroborating their effects.

## METHODS

### Participant

The case study participant was an experienced, male meditator and meditation teacher, aged 52 at the time of data collection. With more than 25 years of experience in ACAM-J, their total estimated practice time amounted to +20,000 hours (based on an estimated daily practice of one hour and one year of retreat with 14 hours of daily practice). While there are different styles of ACAM-J (Sparby and Sacchet, 2024), the case study subject practiced a more sutta-style ACAM-J, reporting use of bodily feelings, breath, and width of attention to enter the form ACAM-J. The participant displayed no neuropsychiatric or cognitive impairment diagnoses as evaluated by the Mini-International Neuropsychiatric Interview (Sheehan et al., 1998) and Mini-Mental State Examination (Folstein et al., 1975) before neuroimaging. The subject provided informed consent, and the study was approved by the Mass General Brigham Institutional Review Board (IRB).

### Experimental Design

#### Advanced concentrative absorption meditation (ACAM-J)

This study employed an intensively sampled case study neurophenomenological design. EEG data were collected throughout 5 days, totaling 29 ACAM-J runs. For each run, the participant progressed without pause through the standard ACAM-J sequence—from the start of the run, termed “access concentration” (AC) on to ACAM-J1 through ACAM-J8. Through this sequence, the subject marked their transition to the next ACAM-J using a button press, excepting ACAM-J6 through ACAM-J8. They did not indicate the later ACAM-J transitions with the button press, as this would have disrupted the deep meditative states and natural flow of their practice. We report the collective period as ACAM-J6-8. Notably, this methodology has demonstrated good within-subject reliability of fMRI brain responses associated with ACAM-J in our previous study (Ganesan et al., 2024). The intensive case-study design enables the collection of high-quality data from a rare, advanced practitioner, leveraging the strength of repeated measurement. In studying advanced meditative phenomena, this approach is particularly powerful, as it reduces the risk of obscuring unique, subject-specific effects and allows for subsequent group sample studies to corroborate the foundational case-study effects, as exemplified by Trautwein et al. (2024) building on the single-case study by Dor-Ziderman et al. (2016).

#### Non-meditative control tasks

To mitigate the potential bias of an adept meditator’s baseline mediative state confounding a traditional resting-state paradigm (Tang et al., 2012), two control tasks were developed to engage the subject in cognitive processes distinct from meditation. The first control was a counting task, in which the participant was instructed to count backward from 10,000 in decrements of 5 with their eyes closed. The second control was a memory task, in which the participants recalled and mentally narrated events of the past week to themselves day by day. The subject completed two runs of each control task, totaling 16 minutes of data for each non-meditative control (8 minutes per run).

#### Phenomenology

To integrate both phenomenological and neuroscientific aspects of ACAM-J investigation, the subject completed first-person phenomenological ratings after each run using a 0 to 10 Likert-type scale. Ratings were administered immediately after each meditation run in order to limit memory bias. This approach allowed for the systematic assessment of the ACAM-J relevant mental and physiological processes and is described at length in prior publications (Yang et al., 2024a). The phenomenological items included were consistent with the typical experience of ACAM-J as described by long-term practitioners (Brasington, 2015) and in Buddhist texts (Khema, 2022), and have shown meaningful relations to patterns of neural activity in prior studies (Ganesan et al., 2024; Yang et al., 2024a; Chowdhury et al., 2025; Potash et al., 2025; Treves et al., in press). Phenomenological items rated for each ACAM-J state included (1) *stability of attention* (ranging from poor to stable); (2) *width of attention* (ranging from narrow to very wide); (3) *intensity of ACAM-J* (how intense a specific ACAM-J quality was). Robustness of the (4) *phenomenology of sights, sounds, physical sensations, and narrative thought stream* during ACAM-J were rated separately for ACAM-J1-J4 (*early phenomenology*) and ACAM-J5-J8 (*late phenomenology*). Additionally, (5) *grades of formlessness* (not having any form) were rated for the formless ACAM-J5-J8. Finally, the subject rated presence of (6) *bliss*/*joy* (intense pleasant bodily sensations) only in ACAM-J2; (7) *cool bliss* (subtle, pleasant mental experience) only in ACAM-J3; and (8) *equanimity* (neutral, neither pleasant nor unpleasant) only in ACAM-J4. We included only stability of attention and width of attention ratings for the phenomenological analyses presented here, as these were directly related to the neural hypothesis that nonlinear dynamics may mediate selective attention processes (Vinck et al., 2023). Additionally, the participant rated these two items for each of the eight ACAM-J, compared to those that were rated for specific groups of ACAM-J. Descriptive statistics and variability for these phenomenological items are reported in Table S6.

### Electroencephalography (EEG) data acquisition and preprocessing

The participant sat approximately 50 cm in front of a computer monitor within an electrically and acoustically shielded booth for EEG data collection. The EEG cap was positioned following the standard international 10-20 system. Continuous EEG signals were recorded using a customized 96-channel actiCAP system and an actiCHamp amplifier (Brain Products GmbH, Gilching, Germany). Electrode impedances were kept below 5 kΩ, and the ground (GND) electrode was integrated into the cap, located anterior and to the right of Channel 10, corresponding approximately to electrode Fz. Channel 1 (Cz) was used as the online reference during data acquisition. All signals were digitized at a sampling rate of 500 Hz using BrainVision Recorder software (Brain Products).

Offline analyses were conducted with MATLAB-based toolboxes EEGLab (Delorme and Makeig, 2004) and FieldTrip (Oostenveld et al., 2011). Preprocessing was briefly as follows: (1) the data were downsampled to 250 Hz, demeaned, and band-pass filtered between 1 and 45 Hz; (2) data were manually inspected for gross muscle and other artifacts, which were manually removed, as well as any noisy channels, which were repaired via interpolation (replaced with the weighted average of neighboring channels as defined by FieldTrip ‘triangulation’ method); (3) EEG time course was detrended to remove linear trends and then referenced to the average of all channels; (4) Independent Component Analysis (ICA) was performed using the automatic component rejection algorithm to remove nonbrain components labeled with less than 5% brain contributions and non-brain combined probability of more than 80%. For ACAM-J runs, runs were divided into distinct state segments (ACAM-J1, ACAM-J2, ACAM-J3, ACAM-J4, ACAMJ5, ACAM-J6-8) based on the subject’s button press indications. For control runs (8 minutes per run, 2 runs of each control type), runs were divided into segments of 1 min for comparable duration to the ACAM-J segments, which yield a total of 16 segments for each control condition. For WPLI and WSMI calculations (see below), we further epoched each ACAM-J and control segments into 5 s epochs, resulting in the following number of epochs across all sessions: ACAM: AC = 196, ACAM-J1 = 452, ACAM-J2 = 553, ACAM-J3 = 699, ACAM-J4 = 541, ACAM-J5 = 366, ACAM-J6-8 = 903. Controls: Counting = 221, Memory = 173.

### WPLI: weighted phase lag index

The WPLI, a widely used metric of phase coherence in EEG data (Vinck et al., 2011; Canales-Johnson et al., 2021), measures the extent to which phase angle differences between two time series *x*(*t*) and *y*(*t*) are distributed towards positive or negative parts of the imaginary axis in the complex plane (similar to the PLI). The underlying idea is that volume-conducted activity accounts for the greatest proportion of detected 0° or 180° phase differences between signals. Therefore, only phase angle distributions predominantly on the positive or negative side are considered to obtain a conservative estimate for real, non-volume conducted activity. The PLI is the absolute value of the sum of the signs of the imaginary part of the complex cross-spectral density Sxy of two real-valued signals *x*(*t*) and *y*(*t*) at time point or trial t. While PLI is already insensitive to zero-lag interactions, the weighted Phase-Lag Index (Vinck et al., 2011) further addresses potential confounds caused by volume conduction, by scaling contributions of angle differences according to their distance from the real axis, as almost ‘almost-zero-lag’ interactions are considered as noise affecting real zero-lag interactions:

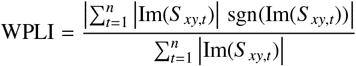

The WPLI is based only on the imaginary component of the cross-spectrum, and thus implies robustness to noise compared to coherence, as uncorrelated noise sources cause an increase in signal power. Here WPLI was computed using the Fieldtrip toolbox (multi-taper method fast Fourier transform, single Hanning taper1, 0.5 Hz frequency resolution). For each frequency range (i.e., 1-3 Hz; 4-7 Hz; 8-14 Hz; 15-29; 30-45 Hz) and epoch, we obtained a single WPLI value after computing and averaging WPLI across all electrode pairs.

### WSMI: weighted symbolic mutual information

We quantified the information sharing between electrodes by calculating the weighted symbolic mutual information (WSMI). This index estimates to which extent two EEG signals exhibit non-random joint (i.e., correlated) fluctuations. Thus, WSMI has been proposed as a measure of neural information sharing (King et al., 2013) and has three main advantages. First, it is a rapid and robust estimate of signals’ entropy (i.e., statistical uncertainty in signal patterns), as it reduces the signal’s length (i.e., dimensionality) by looking for qualitative or ‘symbolic’ patterns of increase or decrease in the signal. Second, it efficiently detects highly nonlinear coupling (i.e., non-proportional relationships between neural signals) between EEG signals, as it has been shown with simulated (Imperatori et al., 2019) and experimental EEG and ECoG data (Canales-Johnson et al., 2020a,b; Olivares et al., 2022; Sitt et al., 2014; King et al., 2013). Third, it rejects spurious correlations between signals that share a common source, thus prioritizing non-trivial pairs of symbols.

We calculated WSMI between each pair of electrodes, for each trial, after transforming the EEG signal into a sequence of discrete symbols defined by ordering of k time samples with a temporal separation between each pair (or τ). The symbolic transformation is determined by a fixed symbol size (κ = 3, i.e., 3 samples represent a symbol) and the variable τ between samples (temporal distance between samples), thus determining the frequency range in which WSMI is estimated (King et al., 2013). The frequency specificity *f* of WSMI is related to k and τ as follows:

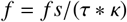

With a *f s* = 250 Hz, as per the above formula, and a *k* size of 3, τ = 64 ms is sensitivity to frequencies ∼1-3 Hz, τ = 32 ms (∼ 4-7 Hz), τ = 16 ms (∼ 8-14 Hz), τ = 8 ms (∼ 15-29 Hz), and τ = 4 ms (∼30-45 Hz). The weights were added to discard the conjunction of identical and opposite-sign symbols, which indicate spurious correlations due to volume conduction. The WSMI (in bits) can be calculated as:

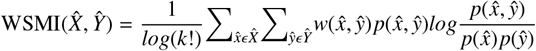

Where x and y are all symbols present in signals *X* and *Y* respectively, *w*(*x, y*) is the weight matrix, and *p*(*x, y*) is the joint probability of co-occurrence of symbol *x* in signal *X* and symbol *y* in signal *Y*. Finally, *p*(*x*) and *p*(*y*) are the probabilities of those symbols in each signal, and *K*! is the number of symbols used to normalize the mutual information (MI) by the signal’s maximal entropy. For each frequency range and epoch, we obtained a single WSMI value after computing and averaging WSMI across all electrode pairs.

### dir-INFO: directed information

To quantify the directed functional connectivity between different EEG signals, we used Directed Information (dir-INFO), also known as Transfer Entropy, an information-theoretic measure of Wiener-Granger causality (Massey et al., 1990). Compared to traditional causality detection methods based on linear models (e.g., Granger causality), dir-INFO is a model-free measure that can detect linear and nonlinear dependencies between brain signals. We took advantage of previous work that made this measure statistically robust when applied to neural data (Ince et al., 2017).

Thus, dir-INFO quantifies functional connectivity by measuring the degree to which the past of a signal *X* predicts the future of another signal *Y*, conditional on the past of *Y*, defined at a specific lag or delay τ:

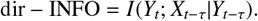

Thus, if there is significant dir-INFO between EEG signal *X* at one time, and EEG signal *Y* at a later time, this shows that signal *X* contains information about the future signal *Y*. Conditioning out the past of signal *Y* ensures the delayed interaction is providing new information over and above that available in the past of signal *X*. For all dir-INFO analyses, we tested delays from 0 ms to 160 ms in steps of 4 ms. For significance testing in each ACAM-J state, we computed a t-test in each time point between feedback and feedforward direction, with a false discovery rate (FDR) correction. dir-INFO was computed on the averaged time series corresponding to two regions of interest (ROI); Front (9 electrodes) and Back (9 electrodes). The corresponding electrodes are depicted in Figure S2).

### Implementation and evaluation of a Naive Bayes classifier for WSMI, WPLI, and phenomenology

#### Model and Training

For each frequency band, a Naive Bayes classifier was selected due to its computational efficiency, robustness to high-dimensional data, and probabilistic output, which is beneficial for understanding prediction confidence. The classifier was trained using features extracted from WPLI and WSMI data, constructing the feature matrix by reshaping the upper triangular part of the connectivity matrices into row vectors, and the corresponding labels Y were assigned based on the ACAM-J. To train and evaluate the model, the data was split into training (70%) and test (30%) subsets. The test data remained completely unseen during training to provide a realistic estimate of the model’s generalization ability. Within the training set, 10-fold cross-validation was performed to ensure robust performance estimation.

#### Performance metrics

The model’s overall accuracy was calculated as the ratio of correctly predicted instances to the total number of cases in the test set. Additionally, a confusion matrix was constructed to visualize the classification performance across different classes, and balanced accuracy was computed to provide a more accurate performance measure in the presence of class imbalance. To gain a detailed understanding of the model’s performance, we calculated precision, recall, and F1-score for each class. Precision was defined as the ratio of true positive predictions to the total predicted positives for a class, recall as the ratio of true positive predictions to the total actual positives for a class, and F1-score as the harmonic mean of precision and recall.

#### Handling class imbalance

Given the class imbalance in the dataset (137 trials for baseline condition, 326 trials for ACAM-J1, 383 trials for ACAM-J2, 479 trials for ACAM-J3, 389 trials for ACAM-J4, 250 trials for ACAM-J5, and 633 trials for ACAM-J6-8), class weights were computed based on the distribution of classes within the training set. These weights were then normalized to create prior probabilities, which were incorporated into the Naive Bayes model to adjust for class imbalance. The class weights were calculated as the inverse frequency of each class in the training set, and these weights were used to scale the prior probabilities accordingly.

#### Phenomenological weighting

Additionally, phenomenological weighting was applied to the features based on the attention dimensions reported in phenomenological reports, specifically stability of attention, and width of attention, ensuring that the feature representation was adjusted according to the underlying phenomena associated with each ACAM-J. The computation of phenomenological weights involved a systematic approach to standardize and integrate subjective and attention ratings across various ACAM-J. The ‘stability of attention’ scale was selected as the basis for these weights because it was consistently present across all ACAM-J. For each ACAM-J, the median of the intensity measurements was calculated, as the median is less sensitive to outliers and provides a robust measure of central tendency. Next, the computed median values underwent min-max normalization to standardize the range of the values. The normalization process involved subtracting the minimum value of the mean scores from each mean value and then dividing by the range (difference between the maximum and minimum values). A small constant (‘epsilon = 1e-5’) was added to the normalized values to prevent potential issues related to zero values. Finally, the normalized mean values were adjusted and combined as necessary to reflect the specific experimental conditions. In particular, the means for ACAM-J6, ACAM-J7, and ACAM-J8 were averaged to produce a single combined mean value. This was necessary because the subject did not mark their transition from ACAM-J6 to ACAM-J8 with a button press, as this would have disrupted the depth and natural flow of their meditation practice. As such, ACAM-J6-8 were treated as a single condition. The final phenomenological weights were then constructed by appending this combined mean to the array of normalized means for ACAM-J1-J5.

These phenomenological weights were subsequently applied in the analysis to adjust for subjective intensity differences between the ACAM-J, thereby enhancing the accuracy and interpretability of the Naive Bayes classifier’s performance across different experimental conditions.

To determine if there was a significant difference in model performance with and without phenomenological weighting, we compared the distributions of F1-Scores obtained under both conditions. Given the nature of F1-Scores and their distribution, we used the Mann-Whitney U test, a non-parametric test, to assess the statistical significance of any observed differences.

#### Stability Analysis

The stability of the Naive Bayes model was assessed through permutation tests. In these tests, the labels in the test set were permuted multiple times, and the model’s accuracy was recalculated for each permutation. This process helped in evaluating the model’s sensitivity to variations in the data. The average accuracy and standard deviation across these permutations were reported to provide insight into the model’s robustness. A p-value was calculated to quantify the statistical significance of the observed accuracy.

### Statistics

Statistical analyses were performed using MATLAB (2023a), and JASP statistical software (open source).

## RESULTS

### Nonlinear versus linear connectivity across ACAM-J

We hypothesized that fluctuations in the phenomenology of self-reported states of attention during ACAM-J (e.g., stability of attention) would involve changes in nonlinear connectivity across distributed electrodes. Thus, we computed and compared between ACAM-J and controls an information theoretical metric capable of capturing nonlinear interactions (WSMI; King et al., 2013; Imperatori et al., 2019). We computed WSMI between ACAM-J and controls at different frequency ranges captured by the variable τ (see Materials and Methods) which measures non-oscillatory connectivity. We used a τ of 64 ms (∼1–3 Hz), 32 ms (∼4–7 Hz), 16 ms (∼8–14 Hz), 8 ms (∼15–30 Hz), and 4 ms (∼30–45 Hz). A one-way ANOVA (ANOVA) of all 5 s epochs per ACAM-J and counting control states (See Methods; number of epochs ACAM-J1 = 452, ACAM-J2 = 553, ACAM-J3 = 699, ACAM-J4 = 541, ACAM-J5 = 366, ACAM-J6-8 = 903, Counting = 221 epochs) showed that WSMI distinguished between ACAM-J, and between the counting control task versus ACAM-J in all frequency ranges (τ: 16 ms, 8 ms, and 4 ms) except for the 1-3 Hz range (τ: 32 ms). WSMI was particularly sensitive to differences between ACAM-J in the 8-45 Hz frequency range (τ: 4 ms) (ANOVA statistics reported in Figure 2A; for descriptive statistics see Table S7). Similar results were observed when comparing ACAM-J to the memory control task (statistics reported in Figure S1A).

**Figure 2:**
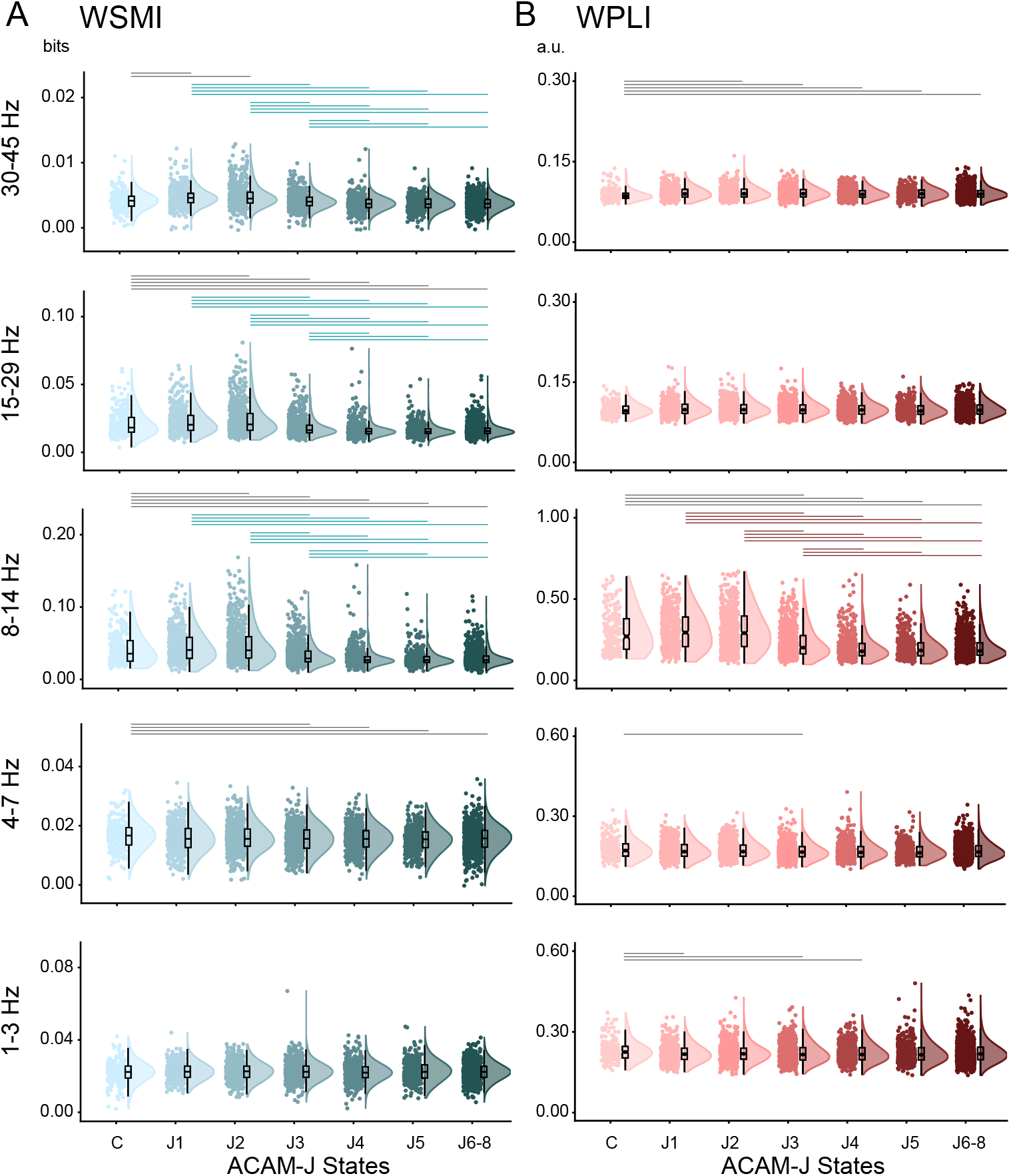
WSMI and WPLI across ACAM-J **(A)** WSMI across ACAM-J and control (Counting) states in different frequency ranges. WSMI (30-45 Hz): One-way analysis of variance (ANOVA) using ‘state’ as a factor with seven levels (Control, and ACAM-J1 to J6-8) revealed a significant main effect of ‘state’ (F_1,3728_ = 50.96; p<0.001). Post-hoc comparisons revealed differences between control and ACAM-J states (C vs J1 and J2; p<0.01), and between ACAM-J (J1 vs J3, J4, J5, J6-8; J2 vs J3, J4, J5, J6-8; J3 vs J4, J5, J6-8; all p<0.01). WSMI (15-29 Hz): main effect (F_1,3728_ = 97.41; p<0.001), with post-hoc differences between control and ACAM-J (C vs J2, J3, J4, J5, J6-8; p<0.01), and between ACAM-J (J1 vs J3, J4, J5, J6-8; J2 vs J3, J4, J5, J6-8; J3 vs J4, J5, J6-8; all p<0.001). WSMI (8-14 Hz): main effect (F_1,3728_ = 108.70; p<0.001), with post-hoc differences between control and ACAM-J (C vs J1, J2, J3, J4, J5, J6-8; p<0.01), and between ACAM-J (J1 vs J3, J4, J5, J6-8; J2 vs J3, J4, J5, J6-8; J3 vs J4, J5, J6-8; all p<0.001). WSMI (4-7 Hz): main effect (F_1,3728_ = 3.04; p = 0.005), with post-hoc differences between control and ACAM-J (C vs J3, J4, J5, J6-8; p<0.02). **(B)** Similarly, a second one-way ANOVA was performed for WPLI values. WPLI (30-45 Hz): main effect of ‘state’ (F_1,3728_ = 5.12; p<0.001). Post-hoc comparisons revealed differences between control and ACAM-J (C vs J2, J3, J4, J5, J6-8; p<0.02). WPLI (8-14 Hz): main effect (F_1,3728_ = 142.58; p<0.001), with post-hoc differences between control and ACAM-J (C vs J3, J4, J5, J6-8; p<0.01), and between ACAM-J (J1 vs J3, J4, J5, J6-8; J2 vs J3, J4, J5, J6-8; J3 vs J4, J5, J6-8; all p<0.001). WPLI (4-7 Hz): main effect (F_1,3728_ = 2.65; p = 0.014), with post-hoc differences between control and ACAM-J (C vs J3; p=0.037). WPLI (1-3 Hz): main effect (F_1,3728_ = 2.13; p = 0.046), with post-hoc differences between control and ACAM-J (C vs J1, J3, J4; p<0.03). Post-hoc tests were Tukey’s corrected for multiple comparisons. Grey horizontal lines represent significant differences between control (Counting) and ACAM-J. Green and brown horizontal lines represent significant differences between ACAM-J for WSMI and WPLI, respectively. Box plots display the sample median alongside the interquartile range. Individual dots represent WSMI (in bits) and WPLI values (arbitrary units; a.u.) for each epoch combining all sessions (see Methods).

Having established that the nonlinear connectivity metric WSMI distinguished self-reported ACAM-J and from control conditions, we compared the results against a well-established linear connectivity metric that captures oscillatory synchronization. To this end, we computed WPLI (weighted Phase Lag Index) (Vinck et al., 2011) in the same frequency ranges as WSMI (Figure 2B). The ANOVA across all epochs showed that, except for the 15-30 Hz range, WPLI distinguished ACAM-J versus the counting control in all other frequency ranges. However, WPLI failed to distinguish between ACAM-J in four out of five frequency ranges (ANOVA statistics reported in Figure 2B; for descriptive statistics see Table S7). Further, this pattern was again observed when comparing ACAM-J to the memory control task (statistics reported in Figure S2B; for descriptive statistics see Table S7).

Taken together, the results supported our hypothesis, showing that WSMI in a broad range of frequencies outperformed WPLI in distinguishing ACAM-J and suggesting that self-reported ACAM-J are associated with changes in nonlinear processing across the brain.

### WSMI outperforms WPLI in decoding ACAM-J

The inferential analyses showing that WSMI outperforms WPLI in distinguishing between ACAM-J raise the question of whether the nonlinear metric WSMI can also significantly decode ACAM-J above chance and, if so, whether decoding accuracy would be higher for WSMI than for WPLI. While inferential analyses allow for the comparison of connectivity means, decoding ACAM-J with the incorporation of both neural (WSMI and WPLI) and phenomenological (ratings of attention) data allows for prediction and comparison of ACAM-J in the presence or absence of self-report subjective ratings. This can inform our understanding of the alignment of subjective experience with underlying patterns in neural dynamics. To answer these questions, we trained a Naive Bayes classifier to decode the meditation states using WSMI and WPLI metrics as features. We first evaluated the classifier performance using WPLI and WSMI in each frequency range as features to decode combined ACAM-J1-J6-8 versus each non-meditative control task, demonstrating that these metrics could successfully distinguish meditation (ACAM-J) and control (counting, memory) states (Table S1). Then, to evaluate our main hypotheses, we evaluated classifier performance, again using WPLI and WSMI in each frequency range, distinguishing specifically within AC and ACAM-J1-J6-8 states (Figure 3). Accuracy, balanced accuracy, precision, recall, and F1-score were computed for each class (see Methods). In both the WPLI and WSMI analyses, the 30-45 Hz range demonstrated superior predictive performance (Figure 3). the WSMI (30-45 Hz) outperformed WPLI, achieving an accuracy of 36.34% and a balanced accuracy of 34.90%. Precision scores ranged from 20.99% (ACAM-J5) to 46.76% (ACAM-J2), recall between 14.17% (ACAM-J5) and 50.19% (ACAM-J6-8), and F1-scores from 16.92% to 46.94%. The ACAM-J6-8 state exhibited the highest performance (F1 = 46.94%), followed by ACAM-J2 (F1 = 39.51%) (Figure 3A). In the case of WPLI (30-45 Hz), the model achieved an overall accuracy of 34.85% and a balanced accuracy of 33.00%. Precision scores for ACAM-J ranged from 19.01% (ACAM-J5) to 52.49% (ACAM-J6-8), with recall scores between 19.17% (ACAM-J5) and 35.58% (ACAM-J6-8). The F1-scores varied between 19.09% (ACAM-J5) and 42.41%, with the highest score observed for ACAM-J6-8 (F1 = 45.51%) (Figure 3B). The results for all other frequency ranges are reported in Table S2) for WSMI, and in Table S3 for WPLI).

**Figure 3:**
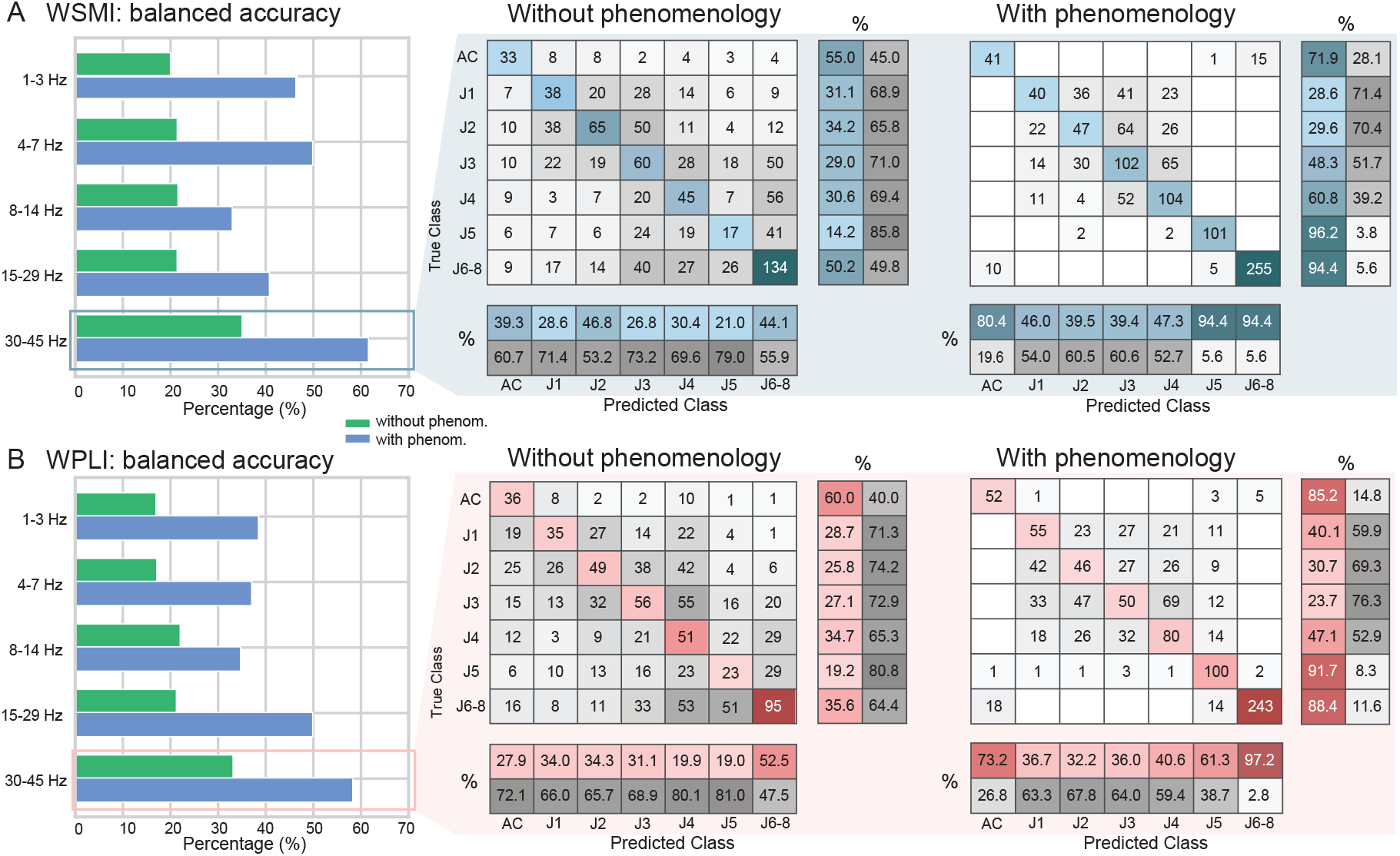
Classification performance of WSMI and WPLI across EEG frequency ranges and ACAM-J meditation states with and without phenomenological data integration. **(A)** WSMI balanced accuracy is shown across the five EEG frequency bands (1–3 Hz, 4–7 Hz, 8–14 Hz, 15–29 Hz, and 30–45 Hz). The bar graph on the left compares model accuracy with (blue) and without (green) phenomenological data inclusion. Confusion matrices on the right show classification performance across baseline control (C) and ACAM-J (J1 to J6-8), indicating true class labels on the y-axis and predicted classes on the x-axis. Matrices depict instances count per cell, with background shading representing classifier performance. The percentages below and at the right side of each matrix represent precision and recall for each state. Precision (shown horizontally at the bottom) indicates the proportion of correctly predicted instances for each state, while recall (displayed vertically along the right side) represents the proportion of true instances that were correctly classified. Higher values in these metrics reflect improved classifier performance in distinguishing between different states. **(B)** WPLI balanced accuracy is displayed following the same structure as **(A)**, with confusion matrices illustrating classification accuracy for models trained with and without phenomenological data. The results demonstrate enhanced classification accuracy, particularly for higher-frequency bands, when phenomenological data are included, suggesting that integrating subjective experience may improve the models’ ability to distinguish between ACAM-J.

To evaluate the statistical significance of decoding performance, we contrasted the WSMI and WPLI results against surrogate data using permutation tests. For WSMI and WPLI metrics, we observed p-values consistently <0.01 (see Methods), indicating that the models’ performances were significantly better than chance. These results show that nonlinear connectivity (WSMI) outperforms linear connectivity (WPLI) in decoding ACAM-J across multiple frequency ranges.

### Integrating self-reported attention with WSMI and WPLI improves decoding of ACAM-J

Can subjective ratings of attention improve the decoding accuracy ACAM-J when combined with the neural connectivity metrics? To answer this question, we used the Naive Bayes classification incorporating the subjective ratings of attention as weights in the model (see Methods). Specifically, we used the subjective ratings of stability of attention, ranging from very poor stability to excellent stabliity, and width of attention, ranging from very narrow to panoramic, as these items were rated by the participant for every ACAM-J state and directly related to our focus on attentional processes during ACAM-J (see Methods). Thus, for each connectivity metric and each frequency range, we repeated the decoding analysis using WSMI and WPLI metrics and incorporated the subjective ratings (i.e., with ‘phenomenological weighting’) (Figure 3A).

When phenomenological weighting was applied, we observed improved performance for both WPLI and WSMI, which again was greater in the 30-45 Hz range. The WPLI achieved an accuracy of 59.95% and a balanced accuracy of 58.13%. Precision scores ranged from 32.17% (ACAM-J2) to 97.20% (ACAM-J6-8), and recall scores between 23.70% (ACAM-J3) and 91.74% (ACAM-J5). The highest F1-scores was observed for ACAM-J6-8 (F1 = 92.57%) (Figure 3B).

In the case of WSMI, decoding performance incorporating phenomenological weighting showed even greater improvement (Figure 3A), with accuracy increasing to 64.47% and balanced accuracy to 61.41%. Precision ranged from 39.38% (ACAM-J3) to 94.39% (ACAM-J6-8), and recall from 28.57% (ACAM-J1) to 96.19% (ACAM-J5). The best performance was observed for ACAM-J5 (F1 = 95.28%). Results integrating WSMI with stability of attention ratings for all other frequency ranges are reported in (Table S4), and for WPLI in (Table S5). Notably, comparable results were obtained when we repeated the decoding analysis integrating ratings of the phenomenological dimension of ‘width of attention’ with the WSMI metrics (Table S4) and with the WPLI metric (Table S5).

We then statistically compared model performance with and without phenomenological weighting. For WPLI, the mean F1-score without phenomenological weighting was 30.63% (S.D. = 7.18%), which increased to 55.26% (S.D. = 23.86%) with phenomenological weighting (Mann-Whitney U = 8.0, p = 0.038). Similarly, with respect to WSMI, the mean F1-score without phenomenological weighting was 33.91% (S.D. = 9.98%), which increased to 61.61% (S.D. = 24.75%) with weighting. A Mann-Whitney U test indicated a statistically significant difference between these conditions (Mann-Whitney U = 8.0, p = 0.038).

These findings highlight the importance of integrating subjective experience data during advanced meditation with neural connectivity metrics that capture attentional states. Furthermore, they underscore the value of focusing on non-oscillatory, nonlinear dynamics in the study of ACAM-J.

### Directed information (dir-INFO)

Having established that the information-connectivity metric WSMI outperforms the spectral-connectivity metric WPLI, and recognizing that ACAM-J can be distinguished through nonlinear dynamics, we subsequently explored variations in information flow between states. To this end, we computed an information-theoretic metric known as Directed Information (dir-INFO) which quantifies directional functional connectivity between neural signals (Ince et al., 2017).

Compared to directional connectivity methods based on linear models (e.g., Granger causality), dir-INFO detects linear and nonlinear interactions between brain signals, and it is statistically robust when applied to neural data as it does not rely on parametric assumptions (Ince et al., 2017; Canales-Johnson et al., 2023). Specifically, dir-INFO quantifies the degree to which the past of a “sender signal” *X* (e.g., EEG traces of posterior electrodes) predicts the future of another “receiver signal” *Y* (e.g., EEG traces of anterior electrodes), conditional on the past of the receiver signal *Y*. Thus, if there is an increase in dir-INFO between EEG signal *X* at one time, and EEG signal *Y* at a later time, this shows that signal *X* contains information about the future signal *Y*. Conditioning out the past of signal *Y* ensures the delayed interaction is providing new information over and above that available in the past of signal *Y* (Figure 1C). We computed dir-INFO across a range of delays (0 to 150 ms) between signals (Figure 4). We assessed neural interactions in both the feedforward (posterior to anterior) and the feedback (anterior to posterior) directions across ACAM-J as they are thought to play a role in processing perceptual contents (Canales-Johnson et al., 2023, 2020b) (see Methods).

**Figure 4:**
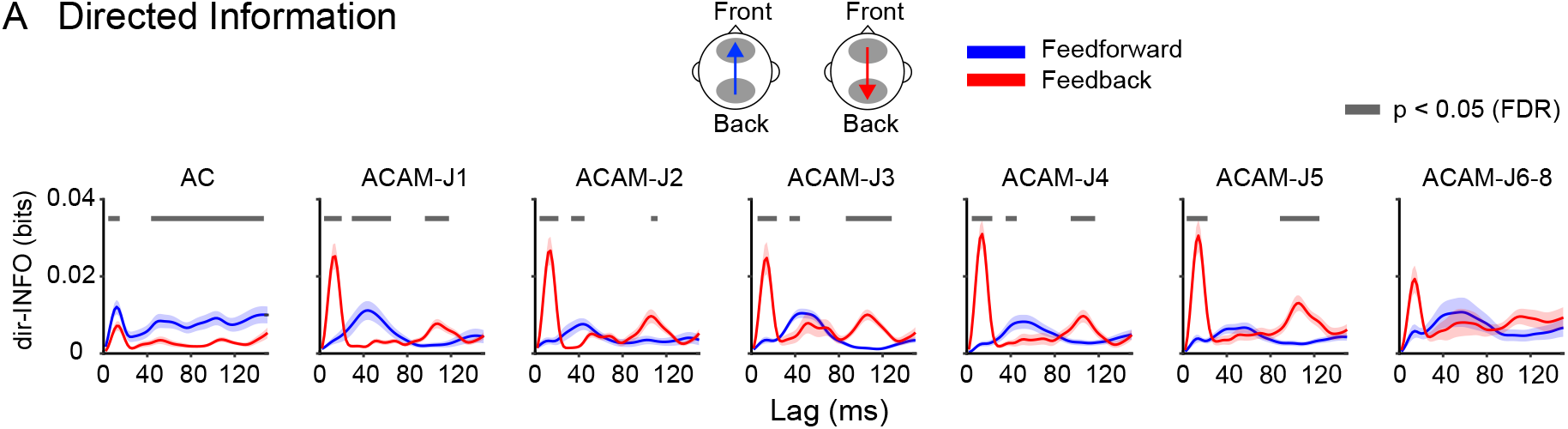
Directed information in ACAM-J **(A)** dir-INFO between the back and front ROIs (feedforward neural information; in blue) and between the front and back ROIs (feedback neural information; in blue) in during Access Concentration (AC) and ACAM-J (J1 to J6-8). Shaded bars represent the standard error of the mean (S.E.M) across epochs. Grey bars represent significant time points after false discovery rate (FDR) correction.

While feedforward information transfer dominated the AC phase before the meditator entered the ACAM-J progression, feedback information transfer was increased in subsequent ACAM-J, suggesting a shift from externally to internally driven perceptual experiences. Notably, the formless ACAM-J showed no difference between feedforward and feedback dir-INFO, suggesting a balance between externally and internally driven neural processing.

## DISCUSSION

This study computed distinct connectivity metrics that capture linear and nonlinear modes of functional brain connectivity. Employing informational theoretic (WSMI and Directed Information) and spectral connectivity metrics (WPLI), we investigated brain connectivity regimes associated with non-oscillatory and oscillatory neural dynamics, respectively. Our results underscored three main findings: (1) WSMI, capturing nonlinear interactions, outperforms WPLI in distinguishing between ACAM-J and control states as well as between ACAM-J states themselves (Figure 2); (2) While our Naive Bayesian classifier distinguished AC and ACAM-J using both WPLI and WSMI connectivity metrics alone, integrating the subject’s phenomenological ratings of the stability of their attention for each ACAM-J as classifier weights increased the model performance (Figure 3); (3) As the meditator moved deeper in the ACAM-J progression, feedforward processes were first reduced and then stabilized, coming to a balance of feedforward versus feedback processes in the later ACAM-J6-8, suggesting a shift from externally to internally driven perceptual experiences (Figure 4).

First, we aimed to investigate nonlinear (WSMI) and linear connectivity (WPLI) during the ACAM-J, specifically how these metrics might be associated with the changing perceptual contents and concentrative processes. Contrary to traditional oscillatory synchronization metrics, nonlinear connectivity captures rich, multi-dimensional dynamics important in pattern extraction and flexible integration of perceptual and cognitive information (Vinck et al., 2023). We hypothesized that the deep absorption, concentration, and filtering of external stimuli during the ACAM-J trajectory would be associated with nonlinear neural interactions distributed across the brain. Indeed, WSMI significantly differed within ACAM-J and between ACAM-J and the counting control in the 30-45 Hz, 15-29 Hz, and 8-14 Hz range, while WPLI only differed within ACAM-J and between ACAM-J and the counting control in the 8-14 Hz range (Figure 2). WSMI generally decreased with deeper states of ACAM-J. This is consistent with prior studies that have reported increased WSMI associated with states of higher arousal (King et al., 2013; Sitt et al., 2014; Imperatori et al., 2019), in contrast to ACAM-J’s increasingly equanimous and stable progression. Additionally, most differences in WSMI of ACAM-J emerged between both non-meditative control tasks and earlier ACAM-J1-J3 compared to ACAM-J4-J6-8. These patterns are coherent with the strong phenomenological differences between the form ACAM-J (ACAM-J1-J4)—-in which the meditator still maintains a point of focus on the ACAM-J object–and the formless ACAM-J (ACAM-J5-J8)—-in which the meditator’s point of focus can no longer be conceptually represented and the awareness of the body falls away (Brahm, 2005; Shankman, 2008; Yang et al., 2024b).

The nonlinear connectivity metric WSMI also outperformed the linear connectivity metric WPLI in Bayesian decoding of ACAM-J (30-45 Hz range), where WSMI model accuracy and balanced accuracy were 36.34% and 34.28%, respectively, while WPLI model accuracy and balanced accuracy were 35.82% and 30.59%, respectively. Both the WSMI and WPLI models’ (30-45 Hz) performance above chance at decoding ACAM-J suggests that both nonlinear and linear interactions may be associated with the cognitive processes supporting ACAM-J. However, WSMI’s stronger distinction of ACAM-J compared to WPLI across both inferential analyses results (Figure 2) and Naive Bayes classifier results (Figure 3) suggest that ACAM-J may be better characterized by nonlinear interactions compared to oscillatory neural connectivity. Crucially, this relates to the hypothesis that self-reported attention and perception modulate more nonlinear than linear connectivity (Vinck et al., 2023; Singer, 2021). ACAM-J are characterized by a progression of states in which the meditator accesses more subtle layers of consciousness as they focus their attention on increasingly stable and refined objects (Yang et al., 2024b). This finding, that WSMI can more effectively capture perceptual changes across the meditation trajectory, highlights the potential significance of non-oscillatory dynamics in these cognitive processes. Nonlinear connectivity’s ability to distinguish meditation-induced altered states of consciousness also aligns with results showing that WSMI distinguishes pathological states of consciousness in vegetative and minimally conscious patients (King et al., 2013; Sitt et al., 2014) and sleep/drowsiness conditions (Imperatori et al., 2019; Canales-Johnson et al., 2020a).

Furthermore, WSMI maintained superior performance in distinguishing ACAM-J compared to WPLI when the subject’s ratings of stability of attention for each ACAM-J were added into the Naive Bayes classifier as phenomenological weights. The boost in performance of both WSMI and WPLI models with phenomenological weighting compared to those without (WSMI mean F1-score increase from 28.41% to 55.26%, p = 0.02; WPLI mean F1-score increase from 33.15% to 61.61%, p = 0.017), underscores the value of taking a neurophenomenological approach in investigating advanced meditative states and other conscious states. Here, we observed that when combined with WSMI, a nonlinear connectivity metric we hypothesize to be more related to modulation of attention and internally-driven perceptual processes, incorporating the subject’s ratings of the stability of attention in each ACAM-J further increased the classifier’s performance in distinguishing ACAM-J.The apparent alignment between subjective ratings of attention stability and nonlinear connectivity metrics supports the notion that non-oscillatory dynamics play a role in modulating the processes associated with subtle shifts in attention and internal perception that define ACAM-J. These results align with a growing body of literature demonstrating how collecting and integrating phenomenological data with neuronal measurements can provide valuable characterizations of neural dynamics underlying complex conscious states (Varela, 1996; Lutz, 2002; Olivares et al., 2015; Ganesan et al., 2024; Timmermann et al., 2023). However, we note that these results are based on an intensively sampled case study dataset, and further validation on independent datasets is necessary to establish broader applicability. Future work should implement similar neurophenomenological decoding models across a wider cohort to support their robustness and generalizability. Mirroring the phenomenological weighting results reported here, in an analysis of regional homogeneity (ReHo) values derived from fMRI data during ACAM-J from the same subject reported in this study, Ganesan et al. (2024) found that incorporation of phenomenological ratings of stability of attention, width of attention, and intensity of ACAM-J increased overall reliability of fMRI responses. It should be noted that in this analysis, we did not exhaust the full phenomenological range of measures to inform the Naive Bayes classifier, but rather we selected phenomenological items to include based on targeted hypotheses of attention-related processes. Although Likert scales are useful for characterizing the relationship between subjective experience and internally driven perception (Lanfranco et al., 2021), future studies should develop a more comprehensive phenomenological characterization of ACAM-J in the context of neural classifications. Moreover, future studies might additionally use subjective measures that provide more granularity than Likert scales, like those implemented in this study design, to capture enhanced temporal dynamics and the richness of subjective experience. For example, retrospective subjective tracings may be beneficial toward providing even stronger links between neural dynamics and temporal dynamics of subjective experience (Lewis-Healey et al., 2024).

Lastly, investigating ACAM-J state-specific patterns in information transfer (dir-INFO) revealed contrasting patterns of feedforward and feedback processes as the meditator progressed deeper in the ACAM-J (Figure 4). In the initial AC phase, feedforward processes appeared to dominate, possibly due to the meditator’s concentration on their breath and bodily feelings that they used to enter the ACAM-J progression. Previously, Yang et al. (2024b) proposed one of the existing frameworks within the meditation research field that may conceptually explain ACAM-J is the many-to-(n)one model of the predictive mind (Laukkonen and Slagter, 2021). This model proposes that focused attention meditation, often practiced when beginning a meditation, may involve an increase in precision-weighting of present moment sensations (i.e., feedforward processes) so as to relatively down-weight distracting predictions that are higher in the neural hierarchy (i.e., feedback processes). In contrast, the model posits that during open monitoring meditation, the practitioner treats all arising signals non-preferentially, and any content of experience is assigned equal, low precision. Additionally, Prest and Berryman (2024) contextualize meditative defabrication within ACAM-J and other deep meditative states using a related active interference framework in which—from ACAM-J1 to ACAM-J8—the meditator experiences a continual decrease in the conceptual complexity of abstraction of phenomenal experience and decreased mental tensing corresponding to a hierarchical level-specific reduction in belief precision. Later in ACAM-J6-8, we observed a relative reduction of the neural system’s reliance on externally-driven information transfer compared to AC and internally-driven information transfer in earlier ACAM-J (ACAM-J1-J5) and instead demonstrated a relative equilibrium of feedforward and feedback processes. This balance of externally-driven and internally-driven information processing may reflect a profound present-moment awareness and interoceptive sensitivity, associated with the deepening concentration and stability of the later ACAM-J (Yang et al., 2024b). In the later ACAM-J6-8, the meditator is increasingly disengaged from external sensations, propagation of prediction errors, or higher level predictions (e.g., thoughts), aligning with the predictive processing and active interference frameworks of Laukkonen and Slagter (2021) and Prest and Berryman (2024). This also aligns with the phenomenology that has been historically reported in relation to the formless ACAM-J more broadly, which are characterized as states of deep equanimity and where sensations as well as narrative and cognitive processing are inhibited (Yang et al., 2024b).

The current findings also align with the hypothesis that transient, nonlinear dynamics characterize changes in perceptual content, whereas stabilization of cognitive states during the late phase of sensory processing relies on oscillatory, linear dynamics (Vinck et al., 2024). ACAM-J illustrate a combination of both change and stability of cognitive states. In ACAM-J, there are shifts in the subtlety of perceptual contents and focus of attention (change) as well as maintenance of overall alertness and sustained attention (stabilization). Indeed, in the current study, both WPLI and WSMI metrics (tracking linear and nonlinear dynamics, respectively) successfully decoded ACAM-J above chance, although WSMI did so with higher accuracy. Future research could further explore the relationship between non-oscillatory and oscillatory mediated sensory processing phases by instructing participants to either (1) maintain or (2) change meditative states. Based on our findings, it could be hypothesized that WPLI would better reflect state maintenance and WSMI would better reflect perceptual content changes.

### Limitations and Generalizability

This study has considerable strengths, including a high volume of data from a single subject during a specific advanced meditation, a rich analysis approach encompassing both oscillatory and non-oscillatory dynamics, and an innovative integration of first-person phenomenology with neural data. However, several limitations of the study should be noted. First, although extensive sampling of a single subject and event has been proven to be an important neuroscientific tool beyond insights that are based on a single or few repeats (Levenson et al., 2012; Poldrack et al., 2015), data from a single subject limits the generalizability of our results. Additionally, compromised button press accuracy during heightened absorption of deeper ACAM-J states (ACAM-J6-8) for the current study subject limits the granularity of results from these later stages. Future research with a diverse group sample of ACAM-J meditators from different traditions and expertise levels is essential for replicating and expanding these findings findings among larger cohorts (Chowdhury et al., 2024; van Lutterveld et al., 2024b). Therefore, while this study offers meaningful contributions in guiding future investigations of nonlinear neural dynamics and advanced meditative states, its findings should be interpreted with caution when extrapolating to broader meditator populations, underscoring the need for further empirical validation across diverse samples.

Overall, we found that while ACAM-J is associated with both linear and nonlinear dynamics, effects related to nonlinear connectivity were stronger. Additionally, nonlinear connectivity data weighted with first-person ratings of stability of attention in each ACAM-J produced the highest accuracy in decoding ACAM-J, which underscores the importance of both nonlinear dynamics and phenomenological reports in the distinction of each ACAM-J. Finally, we observed that late ACAM-J —wherein consciousness is increasingly refined and sensation and narrative processing are inhibited— indicates a balance of externally- and internally-driven information processing, as evidenced by patterns in our analysis of feedforward and feedback processes. These results demonstrate the importance of nonlinear dynamics in distinguishing altered states of consciousness as well as underscore the value of neurophenomenological methods for studying advanced meditation and unique conscious states more broadly. Importantly, the broader neuroscientific characterization of meditation can inform our understanding of consciousness and provide insight into meditation’s beneficial effects on well-being (Abellaneda-Pérez et al., 2024).

## Data Availability

The data and software code that support the findings of this study are available from the corresponding author upon reasonable request, and in alignment with the data sharing policies of the IRB-granting institution (Massachusetts General Hospital).

## Authorship contribution

Conceptualization and methodology: ACJ and MDS. Investigation and data collection: RMP and MDS. Data analysis: SVM, ME and ACJ. Visualization: SVM, ME and ACJ. Writing and editing: RMP, SVM, ME, ACJ and MDS. Supervision: ACJ and MDS. Funding acquisition: MDS.

## Acknowledgements

ACJ is funded by an ANID/FONDECYT Regular (1240899) and ANID/FONDECYT Regular (1251273) research grants. MDS and the Meditation Research Program are supported by the National Institute of Mental Health (R01MH125850), Dimension Giving Fund, Ad Astra Chandaria Foundation, Tan Teo Charitable Foundation, and other individual donors.

---

**Figure S1:**
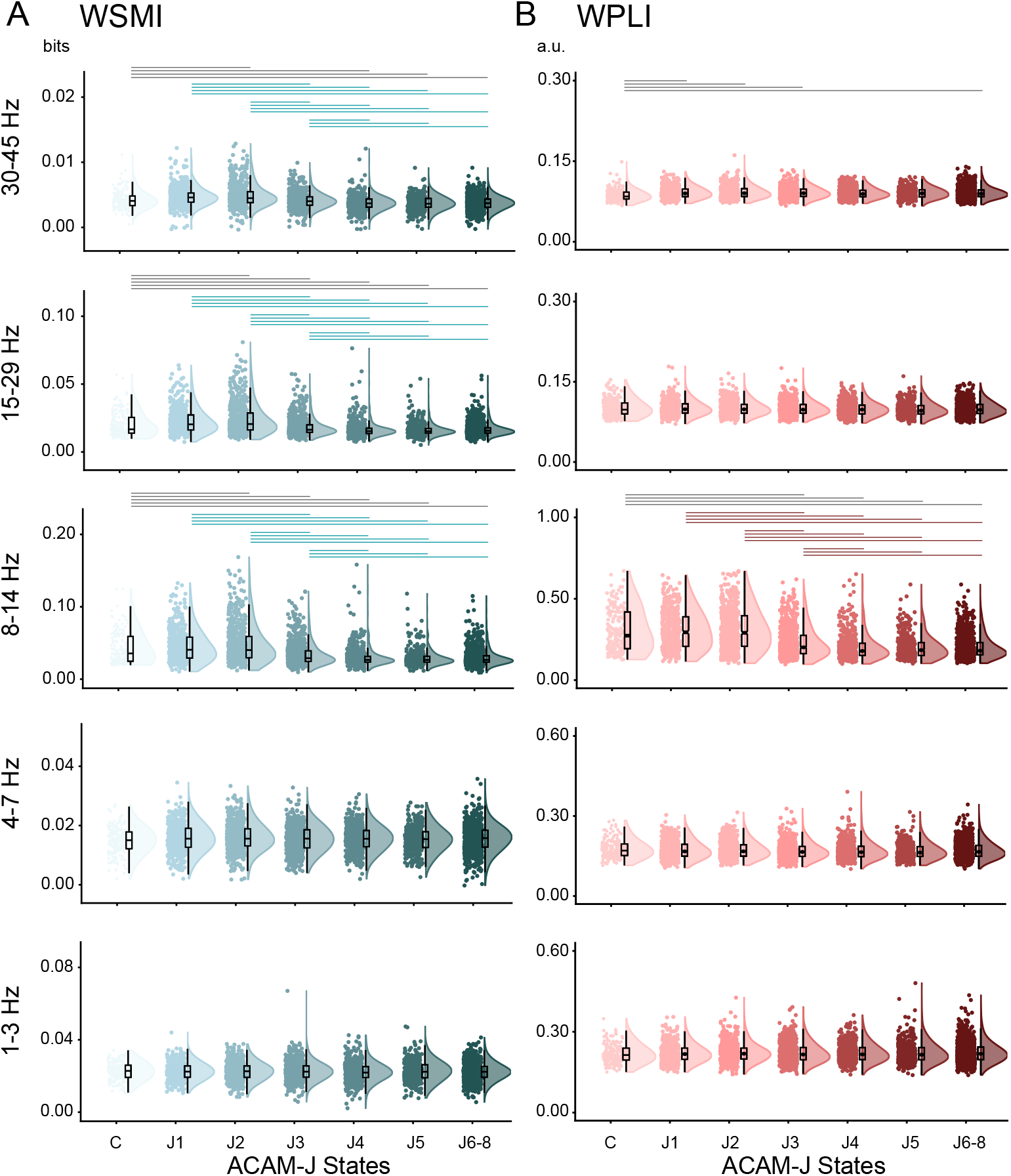
WSMI and WPLI across ACAM-J **(A)** WSMI across ACAM-J and control (Memory) states in different frequency ranges. WSMI (30-45 Hz): One-way analysis of variance (ANOVA) using ‘state’ as a factor with seven levels (Control, and ACAM-J1 to J6-8) revealed a significant main effect of ‘state’ (F_1,3728_ = 53.58; p<0.001). Post-hoc comparisons revealed differences between control and ACAM-J states (C vs J2, J4, J5, J6-8; p<0.01), and between ACAM-J (J1 vs J3, J4, J5, J6-8; J2 vs J3, J4, J5, J6-8; J3 vs J4, J5, J6-8; all p<0.01). WSMI (15-29 Hz): main effect (F_1,3728_ = 99.09; p<0.001), with post-hoc differences between control and ACAM-J (C vs J2, J3, J4, J5, J6-8; p<0.01), and between ACAM-J (J1 vs J3, J4, J5, J6-8; J2 vs J3, J4, J5, J6-8; J3 vs J4, J5, J6-8; all p<0.001). WSMI (8-14 Hz): main effect (F_1,3728_ = 110.03; p<0.001), with post-hoc differences between control and ACAM-J (C vs J2, J3, J4, J5, J6-8; p<0.01), and between ACAM-J (J1 vs J3, J4, J5, J6-8; J2 vs J3, J4, J5, J6-8; J3 vs J4, J5, J6-8; all p<0.001). **(B)** Similarly, a second one-way ANOVA was performed for WPLI values. WPLI (30-45 Hz): main effect of ‘state’ (F_1,3728_ = 3.24; p = 0.002). Post-hoc comparisons revealed differences between control and ACAM-J states (C vs J1, J2, J3, J6-8; p<0.05). WPLI (8-14 Hz): main effect (F_1,3728_ = 140.79; p<0.001), with post-hoc differences between control and ACAM-J (C vs J3, J4, J5, J6-8; p<0.01), and between ACAM-J (J1 vs J3, J4, J5, J6-8; J2 vs J3, J4, J5, J6-8; J3 vs J4, J5, J6-8; all p<0.001). Post-hoc tests were Tukey’s corrected for multiple comparisons. Grey horizontal lines represent significant differences between control (Memory) and ACAM-J. Green and brown horizontal lines represent significant differences between ACAM-J for WSMI and WPLI, respectively. Box plots display the sample median alongside the interquartile range. Individual dots represent WSMI (in bits) and WPLI values (arbitrary units; a.u.) for each epoch combining all sessions (see Methods).

**Figure S2:**
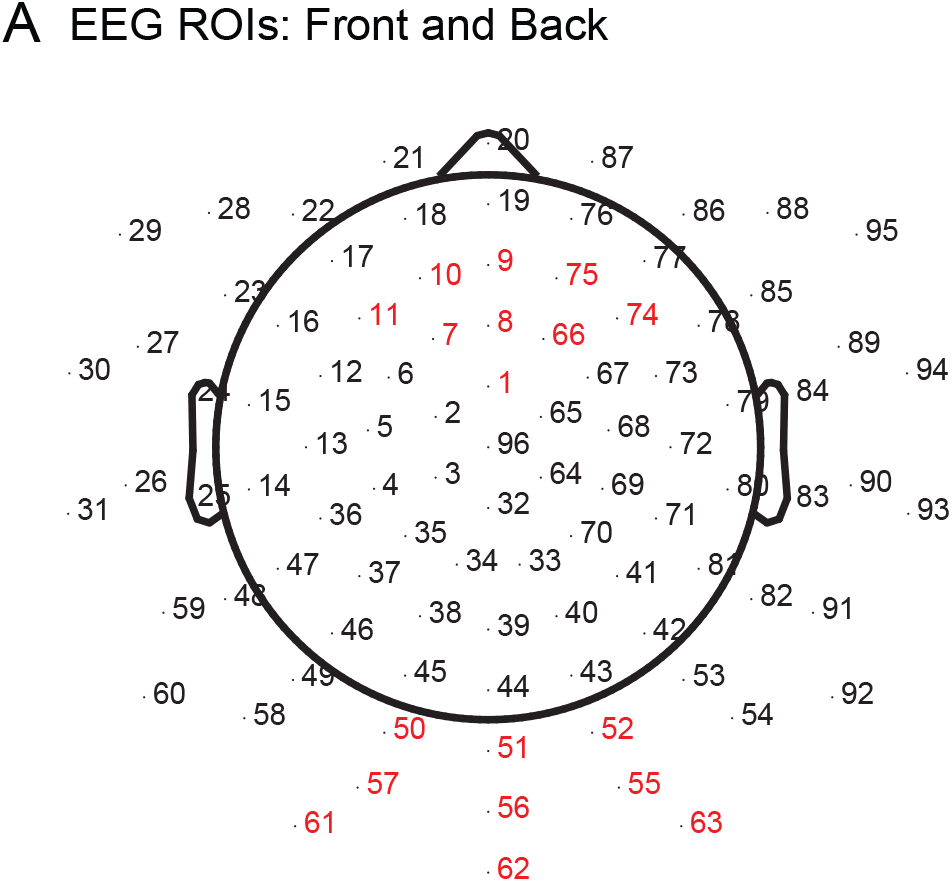
Regions of interest (ROI) for computing dir-INFO in the EEG recordings **(A)** Frontal ROI (Front) is formed by electrodes 1, 7, 8, 9, 10, 11, 66, 74 and 75. Posterior ROI (Back) is formed by electrodes 50, 51, 52, 55, 56, 57, 61, 62 and 63.

**Table S1:**
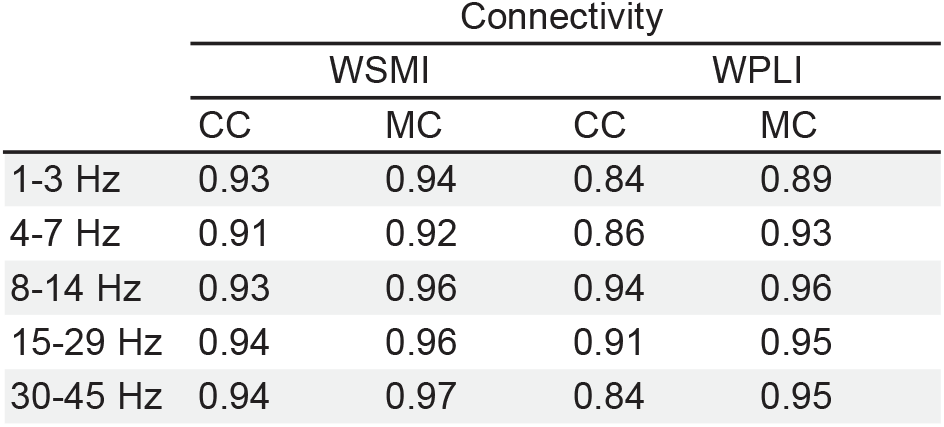
Binary classification accuracies for distinguishing control conditions, Counting Control (CC) and Memory Control (MC), from all ACAM-J (J1 to J6–8) using WSMI and WPLI connectivity metrics. The table presents decoding performance across five EEG frequency bands (1–3 Hz, 4–7 Hz, 8–14 Hz, 15–29 Hz, and 30–45 Hz). Results reflect the binary classification accuracy for each control condition, highlighting the effectiveness of WSMI and WPLI metrics in differentiating control states from meditative states within the specified frequency ranges.

|  | Connectivity |  |  |  |
| --- | --- | --- | --- | --- |
|  | WSMI |  | WPLI |  |
|  | CC | MC | CC | MC |
| 1-3 Hz | 0.93 | 0.94 | 0.84 | 0.89 |
| 4-7 Hz | 0.91 | 0.92 | 0.86 | 0.93 |
| 8-14 Hz | 0.93 | 0.96 | 0.94 | 0.96 |
| 15-29 Hz | 0.94 | 0.96 | 0.91 | 0.95 |
| 30-45 Hz | 0.94 | 0.97 | 0.84 | 0.95 |

**Table S2:** Decoding ACAM-J using WSMI without phenomenological weighting. The table reports overall accuracy, balanced accuracy, and performance metrics (Precision, Recall, and F1-score) across five EEG frequency bands (1–3 Hz, 4–7 Hz, 8–14 Hz, 15–29 Hz, and 30–45 Hz). The chance level for the multiclass classifier with 8 classes is 12.5% (1/8). All reported accuracies and balanced accuracies across frequency bands exceed the chance level. Accuracy is highest for the 30–45 Hz band (38.34%), while balanced accuracy shows consistent performance across bands, with a peak at 30–45 Hz (34.90%). Precision, Recall, and F1-score, reported for each ACAM-J state, highlight variability in classification performance across states and frequency bands. Notably, these metrics underscore the classifier’s ability to balance correct identification of each state (Recall) with minimizing false positives (Precision), resulting in high F1-scores for certain states in the higher frequency bands. The results show the ability of WSMI connectivity metric to distinguish Access Concentration (AC) and ACAM-J states, with particularly strong performance in higher frequency bands.

|  | 1-3 Hz | 4-7 Hz | 8 - 14 Hz | 15-29 Hz | 30-45 Hz |
| --- | --- | --- | --- | --- | --- |
| Accuracy | 22.72 | 25.48 | 18.54 | 17.27 | 36.34 |
| Balanced accuracy | 19.87 | 21.28 | 21.44 | 21.26 | 34.90 |
| (Precision, Recall, F1) |  |  |  |  |  |
| AC | (10.38, 36.67, 16.18) | (18.37, 45.00, 26.09) | (16.67, 6.67, 9.52) | (15.00, 5.00, 7.50) | (39.29, 55.00, 45.83) |
| ACAM-J 1 | (22.92, 18.03, 20.18) | (16.50, 27.05, 20.50) | (25.26, 19.67, 22.12) | (21.21, 17.21, 19.00) | (28.57, 31.15, 29.80) |
| ACAM-J 2 | (20.42, 15.26, 17.47) | (18.81, 10.00, 13.06) | (44.25, 26.32, 33.00) | (44.86, 25.26, 32.32) | (46.76, 34.21, 39.51) |
| ACAM-J 3 | (23.62, 14.49, 17.96) | (21.55, 12.08, 15.48) | (20.41, 9.6, 13.11) | (16.49, 7.73, 10.53) | (26.79, 28.99, 27.84) |
| ACAM-J 4 | (12.70, 10.88, 11.72) | (12.88, 14.29, 13.55) | (9.09, 2.04, 3.33) | (6.82, 2.04, 3.14) | (30.41, 30.61, 30.51) |
| ACAM-J 5 | (16.19, 14.17, 15.11) | (12.85, 19.17, 15.38) | (13.98, 44.17, 21.24) | (18.03, 27.50, 21.78) | (20.99, 14.17, 16.92) |
| ACAM-J 6-8 | (25.90, 29.59, 27.62) | (27.54, 21.35, 24.05) | (29.92, 41.57, 34.80) | (30.37, 64.04, 41.20) | (44.08, 50.19, 46.94) |

**Table S3:** Decoding ACAM-J using WPLI without phenomenological weighting. The table reports overall accuracy, balanced accuracy, and performance metrics (Precision, Recall, and F1-score) across five EEG frequency bands (1–3 Hz, 4–7 Hz, 8–14 Hz, 15–29 Hz, and 30–45 Hz). The chance level for the multiclass classifier with 8 classes is 12.5% (1/8). All reported balanced accuracies across frequency bands exceed the chance level, with the highest performance observed in the 30–45 Hz band (33.00%). Standard accuracy also exceeds the chance level, peaking at 34.85% for the 30–45 Hz band. AC stands for Access Concentration.

|  | 1-3 Hz | 4-7 Hz | 8 - 14 Hz | 15-29 Hz | 30-45 Hz |
| --- | --- | --- | --- | --- | --- |
| Accuracy | 14.43 | 16.22 | 18.94 | 21.68 | 34.85 |
| Balanced accuracy | 16.82 | 16.99 | 21.9 | 21.10 | 33.00 |
| (Precision, Recall, F1) |  |  |  |  |  |
| AC | (12.40, 25.00, 16.57) | (9.17, 18.33, 12.22) | (9.52, 3.33, 4.94) | (16.28, 23.33, 19.18) | (27.91, 60.00, 38.10) |
| ACAM-J 1 | (14.46, 9.84, 11.71) | (12.05, 16.39, 13.89) | (23.89, 35.25, 28.48) | (18.03, 18.03, 18.03) | (33.98, 28.69, 31.11) |
| ACAM-J 2 | (27.68, 16.32, 20.53) | (23.60, 11.05, 15.05) | (47.57, 25.79, 33.45) | (27.12, 25.26, 26.16) | (34.27, 25.79, 29.43) |
| ACAM-J 3 | (21.28, 9.66, 13.29) | (20.46, 25.60, 22.75) | (20.54, 11.11, 14.42) | (20.27, 21.74, 20.98) | (31.11, 27.05, 28.94) |
| ACAM-J 4 | (12.14, 31.97, 17.60) | (17.76, 12.93, 14.96) | (18.37, 6.12, 9.18) | (16.37, 19.05, 17.61) | (19.92, 34.69, 25.31) |
| ACAM-J 5 | (8.11, 10.00, 8.96) | (10.28, 24.17, 14.43) | (11.92, 38.33, 18.18) | (14.81, 13.33, 14.04) | (19.01, 19.17, 19.09) |
| ACAM-J 6-8 | (23.81, 14.98, 18.39) | (31.11, 10.49, 15.69) | (33.97, 33.33, 33.65) | (31.72, 26.97, 29.15) | (52.49, 35.58, 42.41) |

**Table S4:** Classification performance for decoding ACAM-J states (J1 to J6–8) and Access Concentration, integrating WSMI connectivity metrics with phenomenological weighting for stability and width of attention. The table reports overall accuracy, balanced accuracy, and performance metrics (Precision, Recall, and F1-score) across five EEG frequency bands (1–3 Hz, 4–7 Hz, 8–14 Hz, 15–29 Hz, and 30–45 Hz). Two phenomenological dimensions—stability of attention and width of attention—are analyzed independently. For stability of attention, classification accuracy and balanced accuracy improve markedly in higher frequency bands, peaking in the 30–45 Hz band (64.47% and 61.41%, respectively). Precision, recall, and F1-scores are highest for AC and ACAM-J6–8 in the 30–45 Hz band, demonstrating robust classification in these states. For width of attention, a similar trend is observed, with both accuracy (60.02%) and balanced accuracy (65.22%) reaching their highest levels in the 30–45 Hz band. Performance metrics across states indicate strong discriminative power for ACAM-J2 and ACAM-J6–8, with variability in lower frequency bands. These results underscore the role of phenomenological dimensions in improving the classifier’s ability to decode meditative states, particularly in high-frequency EEG bands.

|  | 1-3 Hz | 4-7 Hz | 8-14 Hz | 15-29 Hz | 30-45 Hz |
| --- | --- | --- | --- | --- | --- |
| Stability of attention |  |  |  |  |  |
| Accuracy | 45.32 | 52.54 | 30.44 | 41.63 | 64.47 |
| Balanced accuracy | 46.46 | 49.72 | 32.83 | 40.62 | 61.41 |
| (Precision, Recall, F1) |  |  |  |  |  |
| AC | (33.68, 52.46, 41.03) | (33.33, 52.63, 40.82) | (16.67, 6.15, 8.99) | (17.14, 10.53, 13.04) | (80.39, 71.93, 75.93) |
| ACAM-J 1 | (25.69, 27.41, 26.52) | (28.57, 18.06, 22.13) | (33.75, 20.45, 25.47) | (39.29, 24.81, 30.41) | (45.98, 28.57, 34.24) |
| ACAM-J 2 | (26.09, 24.42, 25.23) | (30.71, 27.74, 29.15) | (41.28, 27.27, 32.85) | (45.45, 23.26, 30.77) | (39.50, 29.56, 33.81) |
| ACAM-J 3 | (33.70, 29.61, 31.52) | (29.94, 25.00, 27.25) | (27.66, 12.50, 17.22) | (21.93, 12.63, 16.03) | (39.38, 48.34, 43.30) |
| ACAM-J 4 | (34.16, 31.98, 33.03) | (35.92, 58.28, 44.44) | (23.49, 22.41, 22.94) | (29.41, 40.94, 34.23) | (47.27, 60.82, 53.20) |
| ACAM-J 5 | (60.13, 87.62, 71.32) | (79.14, 90.91, 84.62) | (15.45, 53.12, 23.94) | (34.78, 87.27, 49.74) | (94.39, 96.19, 95.28) |
| ACAM-J 6-8 | (86.24, 71.76, 78.33) | (89.21, 75.44, 81.75) | (77.52, 87.91, 82.33) | (83.09, 84.93, 84.00) | (94.44, 94.44, 94.44) |
| Width of attention |  |  |  |  |  |
| Accuracy | 48.46 | 53.99 | 45.22 | 42.06 | 60.02 |
| Balanced accuracy | 47.04 | 53.84 | 31.43 | 35.66 | 65.22 |
| (Precision, Recall, F1) |  |  |  |  |  |
| AC | (43.37, 59.02, 50.00) | (48.61, 61.40, 54.26) | (21.50, 66.15, 32.45) | (24.886, 77.19, 37.61) | (72.55, 64.91, 68.52) |
| ACAM-J 1 | (69.52, 54.07, 60.83) | (82.64, 69.44, 75.47) | (47.87, 34.09, 39.82) | (53.76, 37.59, 44.25) | (87.68, 86.43, 87.05) |
| ACAM-J 2 | (74.59, 78.49, 76.49) | (82.82, 87.10, 84.91) | (32.56, 33.94, 33.23) | (43.70, 30.23, 35.74) | (94.30, 93.71, 94.01) |
| ACAM-J 3 | (39.52, 64.08, 48.89) | (50.00, 73.00, 59.35) | (31.15, 9.13, 14.13) | (27.71, 11.62, 16.37) | (70.77, 87.20, 78.13) |
| ACAM-J 4 | (31.01, 23.26, 26.58) | (33.06, 27.15, 29.82) | (30.43, 4.02, 7.11) | (6.25, 1.17, 1.97) | (48.48, 37.43, 42.24) |
| ACAM-J 5 | (21.05, 22.86, 21.92) | (21.00, 17.36, 19.00) | (11.96, 57.29, 19.78) | (27.36, 26.36, 26.85) | (33.72, 27.62, 30.37) |
| ACAM-J 6-8 | (43.11, 27.48, 33.57) | (48.96, 41.40, 44.87) | (40.78, 15.38, 22.34) | (35.39, 65.44, 45.94) | (55.56, 59.26, 57.35) |

**Table S5:** Classification performance for decoding ACAM-J states and Access Concentration (AC) integrating WPLI connectivity metrics with phenomenological weighting for stability and width of attention. The table reports overall accuracy, balanced accuracy, and performance metrics (Precision, Recall, and F1-score) across five EEG frequency bands (1–3 Hz, 4–7 Hz, 8–14 Hz, 15–29 Hz, and 30–45 Hz). For stability of attention, classification accuracy and balanced accuracy improve consistently with increasing frequency, peaking in the 30–45 Hz band (59.95% and 58.13%, respectively). Precision, recall, and F1-scores are highest for AC and ACAM-J6–8 in this band, indicating the classifier’s robust ability to capture these meditative states. For width of attention, accuracy and balanced accuracy also peak in the 30–45 Hz band (54.15% and 56.23%, respectively), with performance metrics highlighting strong decoding for ACAM-J2 and ACAM-J6–8. Performance in lower bands shows greater variability, with reduced precision and recall for certain states. These results demonstrate the importance of high-frequency EEG bands and phenomenological dimensions in decoding meditative states with WPLI connectivity metrics.

|  | 1-3 Hz | 4-7 Hz | 8-14 Hz | 15-29 Hz | 30-45 Hz |
| --- | --- | --- | --- | --- | --- |
| **Stability of attention** |  |  |  |  |  |
| Accuracy | 32.87 | 36.86 | 29.11 | 45.76 | 59.95 |
| Balanced accuracy | 38.33 | 36.91 | 34.57 | 49.72 | 58.13 |
| (Precision, Recall, F1) |  |  |  |  |  |
| AC | (20.69, 31.03, 24.83) | (20.99, 35.42, 26.36) | (12.68, 16.36, 14.29) | (52.24, 52.24, 52.24) | (73.24, 85.25, 78.79) |
| ACAM-J 1 | (19.32, 22.08, 20.61) | (20.77, 27.74, 23.75) | (40.74, 36.07, 38.26) | (23.23, 18.55, 20.63) | (36.67, 40.15, 38.33) |
| ACAM-J 2 | (26.47, 28.12, 27.27) | (21.10, 15.33, 17.76) | (46.46, 35.54, 40.27) | (34.56, 30.72, 32.53) | (32.17, 30.67, 31.40) |
| ACAM-J 3 | (32.82, 19.63, 24.57) | (30.49, 22.12, 25.64) | (36.67, 14.67, 20.95) | (40.13, 28.11, 33.06) | (35.97, 23.70, 28.57) |
| ACAM-J 4 | (17.74, 13.50, 15.33) | (29.09, 19.05, 23.02) | (25.61, 27.81, 26.67) | (30.56, 42.58, 35.58) | (40.61, 47.06, 43.60) |
| ACAM-J 5 | (36.22, 83.53, 50.53) | (30.68, 71.30, 42.90) | (14.11, 31.19, 19.43) | (59.78, 92.24, 72.54) | (61.35, 91.74, 73.53) |
| ACAM-J 6-8 | (84.28, 70.44, 76.74) | (86.51, 67.39, 75.76) | (73.40, 80.35, 76.72) | (89.02, 83.63, 86.24) | (97.20, 88.36, 92.57) |
| **Width of attention** |  |  |  |  |  |
| Accuracy | 39.28 | 35.51 | 36.85 | 46.67 | 54.15 |
| Balanced accuracy | 38.77 | 34.02 | 32.55 | 48.53 | 56.23 |
| (Precision, Recall, F1) |  |  |  |  |  |
| AC | (23.46, 32.76, 27.34) | (16.83, 35.42, 22.82) | (14.70, 74.55, 24.55) | (50.00, 68.66, 57.86) | (53.75, 70.49, 60.99) |
| ACAM-J 1 | (74.42, 62.34, 67.84) | (55.42, 33.58, 41.82) | (27.75, 39.34, 32.54) | (74.47, 56.45, 64.22) | (82.88, 67.15, 74.19) |
| ACAM-J 2 | (54.37, 70.00, 61.20) | (42.57, 57.33, 48.86) | (32.63, 46.39, 38.31) | (69.59, 77.78, 73.46) | (76.19, 85.33, 80.50) |
| ACAM-J 3 | (34.30, 48.40, 40.15) | (32.68, 44.25, 37.59) | (23.21, 5.78, 9.25) | (39.20, 58.53, 46.95) | (46.75, 68.25, 55.49) |
| ACAM-J 4 | (21.24, 14.72, 17.39) | (32.18, 16.67, 21.96) | (26.51, 14.57, 18.80) | (29.25, 27.74, 28.48) | (38.79, 37.65, 38.21) |
| ACAM-J 5 | (12.67, 22.35, 16.17) | (15.57, 17.59, 16.52) | (24.56, 12.84, 16.87) | (20.31, 22.41, 21.31) | (26.04, 22.94, 24.39) |
| ACAM-J 6-8 | (45.60, 20.80, 28.57) | (43.40, 33.33, 37.70) | (42.79, 34.39, 38.13) | (50.32, 28.11, 36.07) | (62.16, 41.82, 50.00) |

**Table S6:** Mean (M), standard deviation (SD), and coefficient of variability (CV) for phenomenology ratings of each ACAM-J. ACAM-J 6-8 (0.022, 0.006) (0.016, 0.006) (0.026, 0.009) (0.015, 0.004) (0.004, 0.001)

| Measure | ACAM-J1 |  | ACAM-J2 |  | ACAM-J3 |  | ACAM-J4 |  | ACAM-J5 |  | ACAM-J6-8 |  |
| --- | --- | --- | --- | --- | --- | --- | --- | --- | --- | --- | --- | --- |
|  | M<br>(SD) | CV | M<br>(SD) | CV | M<br>(SD) | CV | M<br>(SD) | CV | M<br>(SD) | CV | M<br>(SD) | CV |
| Stability of Attention | 8.00<br>(0.45) | 0.06 | 7.97<br>(0.89) | 0.11 | 7.66<br>(1.18) | 0.15 | 7.79<br>(0.92) | 0.12 | 6.86<br>(1.16) | 0.17 | 6.69<br>(1.37) | 0.20 |
| Width of Attention | 3.17<br>(1.86) | 0.59 | 5.28<br>(1.57) | 0.30 | 8.00<br>(0.64) | 0.08 | 9.52<br>(0.62) | 0.07 | 9.86<br>(0.44) | 0.04 | 9.31<br>(0.83) | 0.09 |

**Table S7:** Median and inter-quartile range for wPLI and wSMI values for Counting Control (CC) task, Memory Control (MC) task, and each ACAM-J across five EEG frequency bands (1–3 Hz, 4–7 Hz, 8–14 Hz, 15–29 Hz, and 30–45 Hz).

|  | 1-3 Hz | 4-7 Hz | 8-14 Hz | 15-29 Hz | 30-45 Hz |
| --- | --- | --- | --- | --- | --- |
| **wPLI** |  |  |  |  |  |
| (Median, IQR) |  |  |  |  |  |
| CC | (0.225, 0.044) | (0.171, 0.047) | (0.270, 0.190) | (0.096, 0.014) | (0.086, 0.010) |
| CM | (0.215, 0.039) | (0.167, 0.038) | (0.266, 0.218) | (0.097, 0.019) | (0.086, 0.014) |
| ACAM-J 1 | (0.217, 0.043) | (0.167, 0.044) | (0.294, 0.184) | (0.099, 0.018) | (0.090, 0.015) |
| ACAM-J 2 | (0.219, 0.041) | (0.167, 0.043) | (0.291, 0.190) | (0.099, 0.017) | (0.090, 0.015) |
| ACAM-J 3 | (0.216, 0.047) | (0.165, 0.038) | (0.201, 0.114) | (0.098, 0.017) | (0.090, 0.014) |
| ACAM-J 4 | (0.216, 0.045) | (0.164, 0.040) | (0.179, 0.077) | (0.098, 0.017) | (0.089, 0.013) |
| ACAM-J 5 | (0.215, 0.046) | (0.163, 0.038) | (0.184, 0.082) | (0.096, 0.016) | (0.090, 0.014) |
| ACAM-J 6-8 | (0.218, 0.047) | (0.166, 0.042) | (0.181, 0.078) | (0.097, 0.017) | (0.090, 0.013) |
| **wSMI** |  |  |  |  |  |
| (Median, IQR) |  |  |  |  |  |
| CC | (0.023, 0.007) | (0.017, 0.006) | (0.035, 0.029) | (0.019, 0.011) | (0.004, 0.002) |
| CM | (0.023, 0.007) | (0.015, 0.005) | (0.036, 0.031) | (0.017, 0.013) | (0.004, 0.002) |
| ACAM-J 1 | (0.030, 0.006) | (0.016, 0.006) | (0.040, 0.029) | (0.021, 0.011) | (0.005, 0.001) |
| ACAM-J 2 | (0.022, 0.006) | (0.015, 0.006) | (0.040, 0.030) | (0.021, 0.013) | (0.005, 0.002) |
| ACAM-J 3 | (0.022, 0.006) | (0.015, 0.006) | (0.028, 0.015) | (0.017, 0.006) | (0.004, 0.001) |
| ACAM-J 4 | (0.022, 0.006) | (0.016, 0.006) | (0.026, 0.008) | (0.016, 0.004) | (0.004, 0.001) |
| ACAM-J 5 | (0.022, 0.007) | (0.015, 0.005) | (0.026, 0.008) | (0.016, 0.004) | (0.004, 0.001) |
| ACAM-J 6-8 | (0.022, 0.006) | (0.016, 0.006) | (0.026, 0.009) | (0.015, 0.004) | (0.004, 0.001) |

## Notes

### Competing Interest Statement

The authors have declared no competing interest.

### Summary of Updates

The introduction and discussion were updated after the peer review.

